# Reduction of oligomer size modulates the competition between cluster formation and phase separation of the tumor suppressor SPOP

**DOI:** 10.1101/2023.02.11.528154

**Authors:** Nafiseh Sabri, Matthew J. Cuneo, Melissa R. Marzahn, Jihun Lee, Jill J. Bouchard, Sivaraja Vaithiyalingam, Madeleine B. Borgia, Jeremy Schmit, Tanja Mittag

## Abstract

Phase separation is a ubiquitous process that compartmentalizes many cellular pathways. Given that the same interactions that drive phase separation mediate the formation of complexes below the saturation concentration, the contribution of condensates vs complexes to function is not always clear. Here, we characterized several new cancer-associated mutations of the tumor suppressor Speckle-type POZ protein (SPOP), a substrate recognition subunit of the Cullin3-RING ubiquitin ligase (CRL3), which pointed to a strategy for generating separation-of-function mutations. SPOP self-associates into linear oligomers and interacts with multivalent substrates, and this mediates the formation of condensates. These condensates bear the hallmarks of enzymatic ubiquitination activity. We characterized the effect of mutations in the dimerization domains of SPOP on its linear oligomerization, binding to the substrate DAXX, and phase separation with DAXX. We showed that the mutations reduce SPOP oligomerization and shift the size distribution of SPOP oligomers to smaller sizes. The mutations therefore reduce the binding affinity to DAXX, but enhance the poly-ubiquitination activity of SPOP towards DAXX. This unexpectedly enhanced activity may be explained by enhanced phase separation of DAXX with the SPOP mutants. Our results provide a comparative assessment of the functional role of clusters versus condensates and support a model in which phase separation is an important factor in SPOP function. Our findings also suggest that tuning of linear SPOP self-association could be used by the cell to modulate its activity, and provide insights into the mechanisms underlying hypermorphic SPOP mutations. The characteristics of these cancer-associated SPOP mutations suggest a route for designing separation-of-function mutations in other phase-separating systems.

## Introduction

Phase separation orchestrates spatial and temporal compartmentalization of the cell (1, 2) and influences fundamental processes such as chromatin compartmentalization (3-5), membrane receptor signaling (6, 7), localized translation (8), and the stress response (9-11). Dysregulation of phase separation can cause neurodegenerative diseases and cancer (12, 13). Hence, phase separation is a fundamentally important process used ubiquitously in cells. Yet, important questions remain regarding the extent of the functional role phase separation plays in several processes. Phase separation is mediated by multivalent interactions and leads to the formation of condensates above the saturation concentration, but these same multivalent interactions can also mediate the formation of higher-order oligomers, or so-called “clusters”, in subsaturated solutions (14, 15). If clusters and condensates are formed via the same interactions, how can their relative functional contributions be determined?

We will address this question for the tumor suppressor Speckle-type POZ protein (SPOP), a substrate recognition subunit of the Cullin3-RING ubiquitin ligase (CRL3). SPOP recruits substrates to the CRL3, and these substrates are subsequently poly-ubiquitinated and degraded (16-18). We have previously demonstrated that SPOP forms linear higher-order oligomers via its two dimerization domains, the BTB and BACK domains (19) (Fig. 1A,B). Concentration-dependent oligomerization of SPOP results in an exponential distribution of oligomer sizes, wherein increasing protein concentrations promote the formation of longer and longer oligomers, but long oligomers are always also in equilibrium with short oligomers (Fig. 1C). Given that each SPOP monomer contains a MATH domain that mediates substrate binding, these SPOP oligomers are inherently multivalent for substrates. Certain substrates themselves contain multiple SPOP-binding motifs (20-22) (Fig. 1D), each of which can bind in the substrate-binding cleft of the MATH domain (Fig. 1E). The resulting multivalent interactions encoded by SPOP and its substrates can mediate phase separation and lead to the formation of SPOP/substrate condensates that coexist with a dilute phase (23).

**Figure 1.**
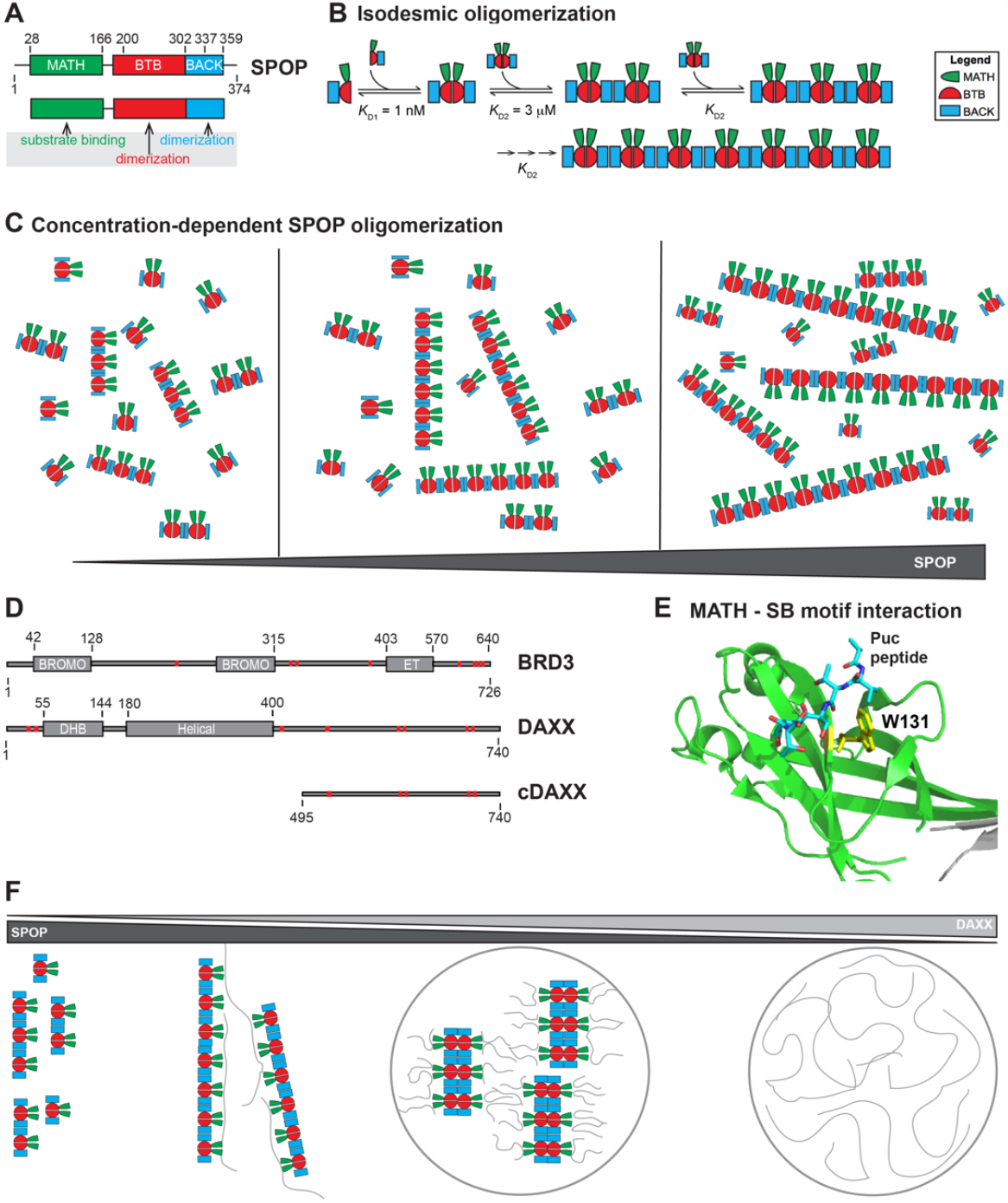
The substrate adaptor SPOP forms higher-order oligomers that are multivalent for substrates. **(A)** SPOP has three functional domains; the MATH domain (green) with the substrate-binding site, and the BTB and BACK domains (red and blue, respectively) that dimerize. **(B)** SPOP undergoes isodesmic oligomerization via its tandem self-association domains: *K*_D1_ and *K*_D2_ are the self-association affinities for the BTB and BACK domains, respectively. *K*_D2_ remains the same independent of oligomer size (19). **(C)** Increasing SPOP concentrations shift the size distribution to longer oligomers, while short and long oligomers are always in equilibrium with each other. **(D)** Schema of the SPOP substrates BRD3 and DAXX. BRD3 has two bromodomains and one extra-terminal (ET) domain. DAXX contains a DAXX helical bundle (DHB) domain and a helical domain. SPOP-binding motifs are depicted as red bars. **(E)** Cartoon model of the MATH domain structure with the canonical SPOP-binding motif from the substrate Puc shown in stick representation. W131 (shown in yellow stick representation) is a residue in the substrate-binding cleft important for substrate binding. **(F)** Schematic representation of possible SPOP, DAXX, and SPOP/DAXX assemblies: SPOP alone forms oligomers (left) (19). At lower molar ratios of DAXX:SPOP, large clusters of SPOP/DAXX are established by stabilization of SPOP oligomers through multivalent DAXX. DAXX can also crosslink oligomers (middle left). At higher molar ratios of DAXX:SPOP, DAXX cannot bind along SPOP oligomers, and SPOP-DAXX brushes are formed, which crosslink via intermolecular DAXX-DAXX interactions and lead to the formation of SPOP/DAXX condensates (middle right). DAXX alone can form condensates (right) (23, 24).

The ability of SPOP to undergo linear oligomerization is required for its sub-cellular localization to nuclear speckles and activity as a substrate recognition subunit of the CRL3 (19). Mutations that disrupt one or both SPOP dimerization interfaces result in constitutive SPOP monomers or dimers and substantial functional defects; the SPOP mutants localize diffusely in the nucleoplasm instead of in nuclear speckles or other nuclear bodies, and they have strongly reduced activity towards substrate poly-ubiquitination (19, 23). But is SPOP oligomerization important because it drives phase separation with substrates (Fig. 1F, middle right panel), or are the higher-order SPOP/substrate oligomers, i.e., the clusters that form below the saturation concentration the functional entities (Figure 1F, middle left panel)? In other words, are phase-separated condensates uniquely suited for function or can clusters perform similar functions? The condensates formed by SPOP and its substrate DAXX have the hallmarks of compartments that are active for SPOP-mediated ubiquitination (23). *In vitro* reconstituted SPOP/DAXX condensates are also active, but so are the clusters that form below the saturation concentration (23). Given that both types of structures are formed by multivalent interactions between SPOP and DAXX, separation-of-function mutations have been difficult to envision. This is the case not only for the SPOP system but in general for multivalent systems that are able to undergo phase separation and also form clusters that may mediate activity.

We previously characterized the network structure of clusters and condensates in the SPOP/DAXX system (24) (Fig. 1F). We demonstrated that these clusters are large SPOP oligomers that are stabilized and cross-linked by the binding of multivalent DAXX. In this system, the resulting clusters are large enough to be light-microscopically observable. At higher DAXX/SPOP molar ratios, multivalent DAXX is not able to stabilize SPOP oligomers and instead leads to the formation of SPOP-DAXX brushes; in these, multivalent DAXX molecules hang off SPOP oligomers. DAXX-DAXX interactions between SPOP-DAXX brushes mediate the formation of condensates above the saturation concentration. Therefore, the network structures (and their underlying interactions) differ between clusters and condensates. These conclusions are also in agreement with theoretical considerations and computational studies of associative polymers, in which cohesive interactions (by so-called stickers, which can make non-covalent physical crosslinks) mediate networking (25, 26), but accompanying density transitions are mediated by the relative insolubility (i.e., poor solvation) of other parts of the molecules (which are typically called spacers) (27, 28). Our recent work also shows that for prion-like low-complexity domains the solubility of the protein molecules (typically encoded in the spacers) as well as their networking ability (via the stickers) together determine the driving force for phase separation (29). Hence, generation of mutations that produce opposite effects on clusters vs condensates should be attainable (15) and would allow for the interrogation of their respective functional contributions.

Here, we assess the molecular mechanism of action of a previously uncharacterized set of cancer-associated SPOP mutations and find that they have opposing effects on the stability of SPOP/substrate clusters and condensates, i.e. they increase the formation of condensates while they reduce the formation of clusters. The mutations target conserved residues in the BTB and BACK domain interfaces and do not appear in healthy individuals but are found in patients with endometrial, skin, and other cancers. We show that the mutations result in a shift in the SPOP oligomer size distribution to smaller sizes relative to WT and reduce binding affinity to multivalent substrates *in vitro*. Paradoxically, the mutations enhance substrate ubiquitination in cells, a surprising observation given the importance of oligomerization for function. We provide a possible molecular explanation for this phenotype by demonstrating that the SPOP interface mutants have a stronger driving force for phase separation together with substrates in vitro. Mutations that differentially affect clusters and phase-separated condensates therefore enable the comparative assessment of these functions. Our results point to the importance of phase separation for SPOP function and suggest a strategy for disentangling the role of clusters vs condensates in other biological processes.

## Results

We and others have previous established the importance of higher-order SPOP oligomerization for its function (19, 23, 30); mutations that completely abrogate dimerization of either or both dimerization interfaces result in altered subcellular localization and loss of ubiquitination activity (19, 23). To probe the impact of more subtle changes to oligomerization on SPOP function, we reviewed cancer genomes for SPOP mutations that may modulate oligomerization. The most prevalent SPOP mutations found in cancer patients are those in the substrate-binding cleft of the MATH domain (Fig. 2A); they reduce substrate binding and turnover of proto-oncogenic substrates and result in prostate cancer oncogenesis (31). By contrast, mutations on the periphery of the MATH domain lead to endometrial cancer (Fig. 2A) and have counterintuitive effects on substrate ubiquitination; some substrates undergo enhanced ubiquitination in their presence while others undergo reduced ubiquitination (32). Additional mutations of unknown significance occur with low prevalence across most of the sequence. However, two uncharacterized mutations in the dimerization domains, R221C and R354H, stood out due to their somewhat higher prevalence as well as their locations (Fig. 2A). The mutated residues are located in the BTB and BACK domain interfaces, respectively, where we expect them to interfere with ion pair bonding across the interface; R221 usually forms a salt bridge with E189, and R354 forms a bridge with E334 (Fig. 2B). R221C was found in 4 patients with skin, mouth, bladder, or large intestine cancers and one additional patient had the mutation R221H; R354H was found in two patients with endometrial or large intestine cancer and 3 additional patients had the mutation R354C. R221 and R354 are conserved across animals (Fig. 2C), and mutation of R221 does not appear as a normal SNP in the population according to the gnomAD database; R354H is seen in one individual. Based on these results, we predicted that the interface mutations reduce linear SPOP oligomerization and that they are potentially pathogenic.

**Figure 2.**
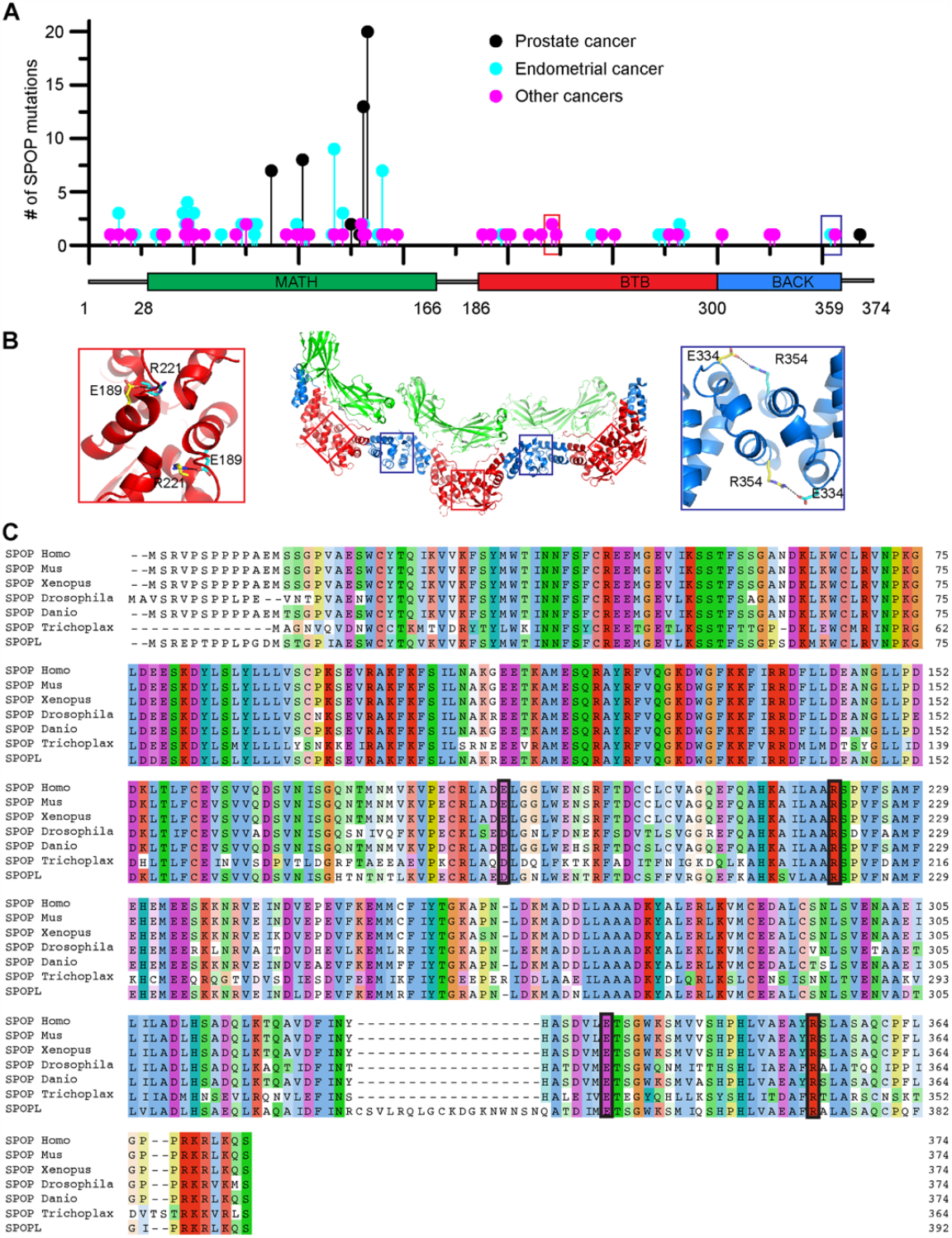
Conserved residues that form contacts across the dimerization interfaces are mutated in cancer patients. **(A)** The lollipop plot shows mutations identified in cancer patients in all three domains of SPOP. Prostate cancer–associated mutations (black) are prevalent in the substrate-binding cleft of the MATH domain. These mutations impair SPOP–substrate interactions, resulting in the stabilization of oncoproteins in prostate cancer patients. Endometrial cancer mutations (blue), which have intermediate frequency, are also clustered in the MATH domain. However, these surprisingly promote ubiquitination of some substrates while inhibiting ubiquitination of others. Among the other SPOP mutations identified (magenta), R221C (red box) and R354H (blue box) are located in the BTB and BACK domains, respectively. Mutation data were collated from cBioPortal (33, 34). **(B)** Model of a SPOP oligomer generated by superimposing the SPOP crystal structures for the BTB dimer (35) and BACK dimer (36). Red and blue boxes show the locations of R221 and R354 in the BTB/BTB and BACK/BACK interfaces, respectively. Both residues form salt bridges with glutamic acid residues across their respective dimerization interfaces. **(C)** Sequence alignment of human SPOP with SPOP homologs in other species and with human SPOP-like protein (SPOPL). The alignment shows evolutionary conservation of R221 and R354 and their corresponding salt bridge partners E189 and E334. Homo, *Homo sapiens*; Mus, *Mus musculus*; Xenopus, *Xenopus laevis*; Drosophila, *Drosophila melanogaster*; Danio, *Danio rerio*; and Trichoplax, *Trichoplax adhaeren*.

### SPOP interface mutants reduce higher-order oligomerization

We used composite-gradient multi-angle light scattering (CG-MALS) experiments to characterize linear oligomerization of SPOP^WT^ and the SPOP mutants (Fig. 3A). We generated a dilution series of SPOP solutions and determined their static light scattering intensities. As expected, SPOP^R221C^ and SPOP^R354H^ showed reduced self-association relative to SPOP^WT^ as evidenced by lower scattering intensities at given protein concentrations (Fig. 3B). To interpret the data, we used an isodesmic oligomerization model in which SPOP dimers (i.e., BTB dimers) self-associate via BACK domain interactions, and the addition of each successive BTB dimer occurs with the same affinity (Fig. 1B). We previously determined the *K*_D_ of BTB dimerization to be ∼1 nM. The *K*_D_ for the isodesmic step was increased from 2.7 µM for SPOP^WT^ to 14.8 µM for SPOP^R221C^ and 26.9 µM for SPOP^R354H^, respectively (Table 1). The mutation R221C is expected to alter the BTB dimerization affinity but we can capture the shift to smaller oligomers with a fit to the same isodesmic model. In conclusion, SPOP interface mutants can form higher-order oligomers, but their size distributions are shifted to smaller sizes (Fig. 3C). The effect of the R221C mutant was more pronounced than that of the R354H mutant. Both interface mutants reduce linear SPOP oligomerization but to different extents.

**Table 1.**

| SPOP variant<br>(residues 28 - 359) | Isodesmic $K_D$ <sup>a</sup> [μM] | $K_D$ for peptide [μM] <sup>b</sup> | $K_D$ for cDAXX [μM] <sup>c</sup> |
| --- | --- | --- | --- |
| WT | $2.7 \pm 0.4$ | $9.8 \pm 0.7$ | $1.2 \pm 0.1$ |
| R221C | $14.8 \pm 2.0$ | $5.9 \pm 0.4$ | $3.2 \pm 0.6$ |
| R354H | $26.9 \pm 2.5$ | $5.1 \pm 0.6$ | $3.0 \pm 0.4$ |
| W131G | N.D. | no binding | N.D. |
| mutBACK | N.D. | $4.3 \pm 0.3$ | N.D. |
| mutBTB | N.D. | $5.7 \pm 0.4$ | N.D. |
| mutBTB-BACK | N.D. | $7.4 \pm 0.4$ | N.D. |

<sup>a</sup> from CG-MAS; mean from at least three replicates ± S.D.
<sup>b</sup> from fluorescence anisotropy assay; mean from at least three replicates ± S.D.
<sup>c</sup> from fluorescence anisotropy assay; mean from at least three replicates ± S.D.
N.D. = not determined

**Figure 3.**
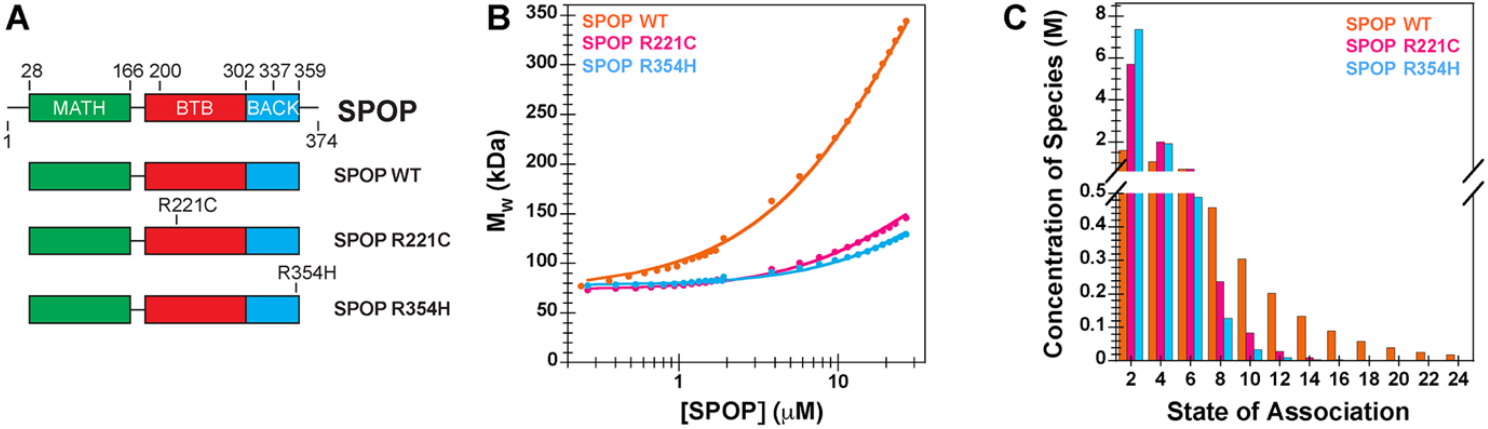
Interface mutations reduce SPOP self-association and oligomer size. **(A)** Cartoon schematics of SPOP^WT^, SPOP^R221C^, and SPOP^R354H^. The experiments with purified protein used constructs comprising residues 28-359; experiments in cells used the full-length protein. **(B)** Experimental weight-average molar mass (M_w_) from CG-MALS (shown in circles) for SPOP WT and the interface mutants were fitted to an isodesmic self-association model in which SPOP dimers are the self-associating unit (solid lines). The largest SPOP oligomer taken into account was an undecamer of SPOP dimers [(SPOP_2_)_11_]. **(C)** Graphical representation of the SPOP concentration of each oligomeric species within a 10 µM solution of SPOP WT or the interface mutants.

### SPOP interface mutants have enhanced ubiquitination activity

To test whether the SPOP interface mutants had functional defects, we established inducible expression systems for SPOP WT and the SPOP mutants in T-REx cells wherein SPOP was expressed at relatively low levels (Fig. S1), and we performed in-cell ubiquitination assays with DAXX as the substrate. While the prostate cancer mutant SPOP^W131G^ had only weak activity as expected, the interface mutants SPOP^R221C^ and SPOP^R354H^ poly-ubiquitinated DAXX more strongly than SPOP^WT^ (Fig. 4A). As a control, we examined the effect of a SPOP mutant harboring the mutation Y353E (resulting in SPOP^mutBACK^) (36), a mutation we previously used to completely abrogate BACK domain dimerization (19, 23). We confirmed that SPOP^mutBACK^ has low ubiquitination activity towards DAXX, which is in agreement with our previous observations that linear higher-order SPOP oligomerization is required for full function (Fig. 4B).

**Figure 4.**
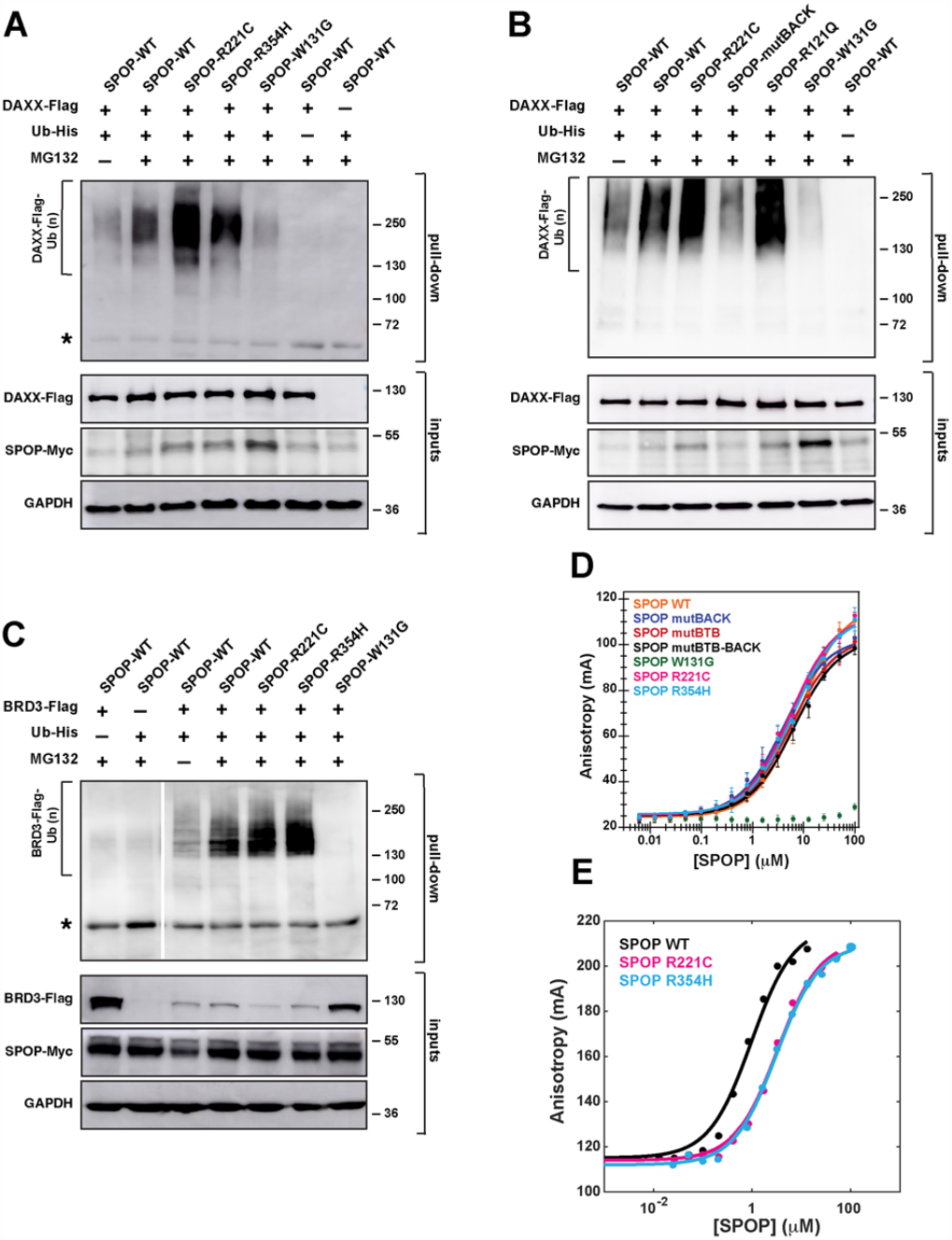
SPOP interface mutations enhance ubiquitination but reduce substrate binding. **(A)** SPOP^R221C^ and SPOP^R354H^ enhance DAXX polyubiquitination. Representative immunoblot showing ubiquitination in T-REx cells transfected to express SPOP-Myc, DAXX-Flag, and His-tagged ubiquitin. Twenty-four hours post-transfection, cells were incubated with DMSO or 20 μM MG132 for 4–5 hours. The cells were lysed and the resulting lysates were used in His-tag pull-down assays, using nickel-NTA beads under denaturing conditions. The samples were then run on SDS–PAGE gels, and the gels were immunoblotted with an anti-Flag antibody. Protein input was verified using antibodies for Myc, Flag, and GAPDH (loading control). **(B)** Similar to the interface mutant R221C, the endometrial cancer mutant R121Q promotes SPOP polyubiquitination activity. The opposite effect is observed for mutBACK, the SPOP mutant that completely disrupts BACK domain dimerization. Experimental conditions as in (A). **(C)** SPOP^R221C^ and SPOP^R354H^ enhance BRD3 polyubiquitination, indicating that gain-of-function for the interface mutants is not limited to the substrate DAXX. Experimental conditions as in (A), except that BRD3-Flag is expressed as the substrate instead of DAXX-Flag. Asterisks in panels A and C indicate a cross-reacting band. **(D)** SPOP affinity for its canonical binding motif is not altered for the SPOP^R221C^ and SPOP^R354H^ mutants relative to SPOP^WT^, but it is reduced in SPOP^W131G^, a prostate cancer mutant that abrogates substrate binding. The affinities were measured using a fluorescence anisotropy binding assay with a fluorescently labeled peptide that was derived from a SPOP-binding motif from the substrate Puckered. **(E)** The binding affinity for SPOP^R221C^ and SPOP^R354H^ to the multivalent substrate is slightly decreased. Affinities were measured using a fluorescence anisotropy binding assay with fluorescently labeled cDAXX.

These observations raised multiple questions: (1) If higher-order oligomerization is required for SPOP activity, why does reduction in oligomerization enhance activity? and (2) Why is enhanced activity of SPOP oncogenic if prostate cancer–causing SPOP mutations reduce substrate ubiquitination? SPOP mutations on the periphery of the MATH domain (including mutant R121Q) that were identified in patients with endometrial cancer enhance turnover of some substrates (32). Given that the mutation R354H was identified in a patient with endometrial cancer, we hypothesized that mutations in different regions of SPOP may have similar effects on substrate turnover and therefore result in similar malignancies. Indeed, the endometrial cancer mutant SPOP^R121Q^ mediated higher poly-ubiquitination activity towards DAXX than SPOP^WT^, mirroring the activity of the SPOP interface mutants (Fig. 4B).

Given the similar activity of SPOP interface and MATH domain mutants associated with endometrial cancer, we tested the activity of the interface mutants towards BRD3, which is a substrate that is ubiquitinated more effectively by SPOP harboring MATH domain mutations found in endometrial cancer patients (32). Indeed, BRD3 ubiquitination was also enhanced when we used the SPOP interface mutants (Fig. 4C). Our functional data suggest that several different types of endometrial cancer mutations enhance SPOP activity towards a set of substrates, though the underlying mechanisms are unknown.

### SPOP interface mutations reduce binding to multivalent substrates

We next sought to understand the mechanism underlying the increase in substrate ubiquitination observed for the SPOP interface mutants. We did not expect a defect in the binding affinity to individual SPOP-binding (SB) motifs given that the MATH domain is intact. Indeed, the binding affinities to the canonical SB motif found in the substrate Puc (35) were identical within error between WT and the interface mutants, and no binding was observed for the binding-incompetent SPOP^W131G^ mutant, as expected (Fig. 4D). Given that the oligomerization state of SPOP may affect the binding of a multivalent substrate, we next investigated the binding affinities between the C-terminal intrinsically disordered region (IDR) of DAXX (cDAXX) and the SPOP mutants. cDAXX has 5 SB motifs (Fig. 1D) (23). As expected, cDAXX bound slightly more tightly to SPOP^WT^ than to the interface mutants (Fig. 4E), presumably due to enhanced avidity from the larger WT oligomers. Thus, a reasonable expectation from binding assays would be that SPOP interface mutants would likely have decreased ubiquitination activity towards multivalent substrates, not enhanced activity.

### SPOP interface mutants enhance phase separation with substrates

To understand the source of the unexpected observation that SPOP interface mutants mediate increased ubiquitination activity, we investigated whether phase separation and discrete binding are affected differently by the mutations. We previously showed that transient expression of SPOP and DAXX results in their colocalization to phase-separated SPOP/DAXX bodies in cells. These bodies also recruited other subunits of the CRL3 and had the hallmarks of an active compartment for SPOP-mediated ubiquitination of DAXX (23).

Thus, we tested the ability of SPOP WT and the interface mutants to undergo phase separation together with cDAXX *in vitro*. We observed condensates containing SPOP WT and cDAXX, and these have the ability to fuse and wet surfaces as demonstrated previously (23) (Fig. 5A). As described previously, different molar ratios of SPOP and cDAXX give rise to different types of assemblies; condensates form at high cDAXX:SPOP molar ratios, and clusters (or filamentous assemblies or gels) form at low cDAXX:SPOP molar ratios (Fig. 1F) (23, 24). Modeling studies showed that the underlying interactions differ (24). Clusters arise when multivalent substrates stabilize SPOP oligomers and crosslink them. By contrast, condensate formation is mediated by DAXX-DAXX interactions between SPOP-DAXX brushes (24).

**Figure 5.**
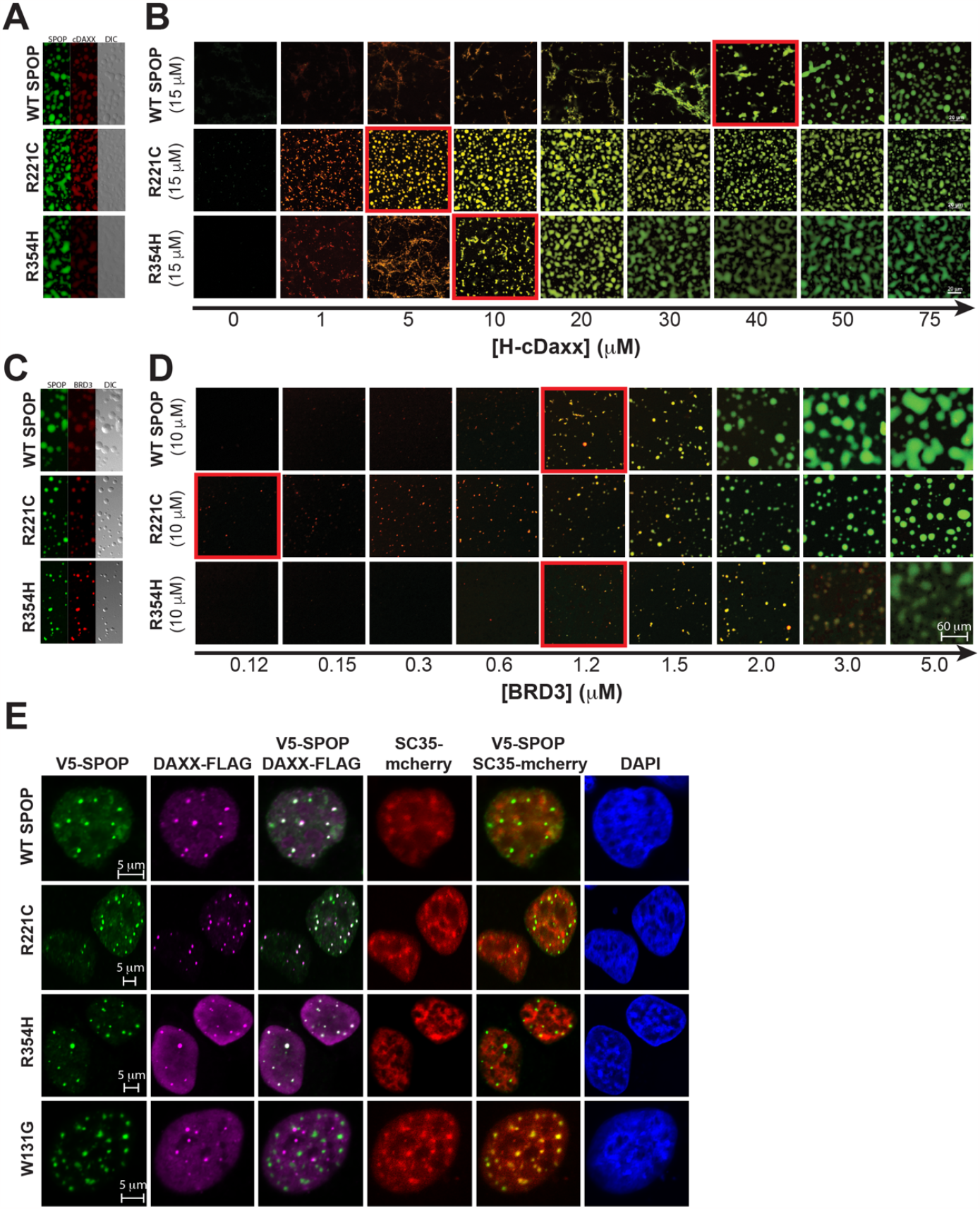
SPOP interface mutants enhance phase separation with substrates. **(A)** Two-channel confocal fluorescence and DIC microscopy images show that SPOP^WT^, SPOP^R221C^, and SPOP^R354H^ each colocalize with cDAXX in SPOP/cDAXX condensates. **(B)** The phase boundary between SPOP/cDAXX clusters and condensates is shifted for the SPOP proteins with interface mutations. Confocal fluorescence microscopy images of SPOP^WT^ or an interface mutant (green) as a function of cDAXX (red) concentration. The boundary between clusters and condensates is indicated by a red outline. All samples contain 10% w/v ficoll 70, 500 nM ORG-SPOP, and/or 500 nM Rhodamine-cDAXX. **(C)** Two-channel confocal fluorescence and DIC microscopy images show that SPOP^WT^, SPOP^R221C^, and SPOP^R354H^ each colocalize with BRD3 in SPOP/BRD3 condensates. **(D)** The phase boundary between SPOP/BRD3 clusters and condensates is shifted for the SPOP proteins with interface mutations. Confocal fluorescence microscopy images of SPOP^WT^ or an interface mutant (green) as a function of BRD3 (red) concentration. The boundary between clusters and condensates is indicated by a red outline. All samples contain 200 nM ORG-SPOP and/or 100 nM Rhodamine-BRD3. **(E)** Representative fluorescence confocal images of HeLa cells expressing V5-SPOP, DAXX-Flag, and SC35-mCherry constructs (the latter to mark nuclear speckles). Cells were transfected with the indicated plasmids. Twenty-four hours post-transfection, cells were fixed and immuno-stained using antibodies against V5 (green) and Flag (magenta). DAPI (blue) marks nuclear DNA.

When we titrated increasing concentrations of cDAXX into SPOP^WT^ solutions, we observed a progression from the formation of clusters at low molar ratios to the formation of condensates at higher molar ratios (Fig. 5B). Condensate formation was demarcated by a saturation concentration. Similar titrations of cDAXX into a solution of SPOP^R221C^ resulted in a lower saturation concentration, i.e., condensate formation at lower cDAXX concentrations (Fig. 5B). SPOP^R354H^ also had a lower saturation concentration but the effect was smaller. These data indicate that the SPOP interface mutants binds to cDAXX more weakly in discrete complexes but engages cDAXX more effectively via phase separation. Similar results were also obtained using BRD3 (Fig. 5C,D).

To investigate whether the SPOP interface mutants can also readily phase separate with DAXX in cells, we determined the cellular localization of the proteins. The SPOP^WT^ and the SPOP interface mutants co-localized with DAXX in SPOP-DAXX bodies (Fig. 5E). By contrast, SPOP^W131G^ did not colocalize with DAXX and was instead localized to nuclear speckles, in agreement with our previous results (23). Under these conditions, DAXX also remained partially localized to PML bodies (Fig. S2), as expected (18, 37, 38). Hence, we conclude that the enhanced ability of the SPOP interface mutants to undergo phase separation with substrates provides one explanation for the increase in ubiquitination activity observed for DAXX and BRD3 in the presence of these mutants. Our attempts to quantify phase diagrams in cells were inconclusive, likely because of the multitude of SPOP substrates in cells that can all contribute to phase separation and vary in levels between cells.

## Discussion

Many different SPOP mutations have been found in cancer genomes (39, 40), and SPOP is regarded as an important tumor suppressor across cancers (41). However, so far only prostate cancer–driving mutations for residues in the substrate binding site are understood at a molecular level; they result in reduced substrate binding (31, 42). Here, we show that two mutations in the dimerization interfaces of the BTB and BACK domains weaken higher-order SPOP oligomerization and shift the size distribution of SPOP oligomers to smaller sizes. While this weakens the binding of multivalent substrates due to reduced avidity, the poly-ubiquitination activity of these mutants towards substrates is higher than that of SPOP^WT^, an initially counterintuitive finding. This apparent contradiction is resolved by the realization that the mutations increase the driving force for phase separation towards substrates, i.e., substrates form condensates with mutant SPOP at lower concentrations. The implication is that a larger fraction of substrate enters the active dense phase and is thus turned over. In addition to revealing biophysical mechanisms that regulate SPOP activity, our results yield a comparative assessment of the functional role of clusters vs condensates and support a model in which phase separation plays an important role in SPOP function.

Why does weakened linear SPOP oligomerization reduce the substrate saturation concentration for phase separation with SPOP? Our previous combined theoretical and experimental characterization of the SPOP/DAXX system provides clues (24). We observed that DAXX has an intrinsic but weak driving force for phase separation. SPOP/DAXX brushes phase separate via these same DAXX-DAXX interactions, but the driving force is substantially higher than DAXX phase separation alone due to the additive effect of tethering multiple DAXX molecules to the SPOP “hub”. However, our data showed that the SPOP/DAXX dense phase has a higher density than the DAXX dense phase (24). Since the pure DAXX dense phase has the optimal density for DAXX, the increased density caused by the SPOP hub must result in an energetic penalty. This penalty will reduce the driving force for DAXX-mediated phase separation below what it could be without this penalty. By contrast, the shorter SPOP oligomers formed by the interface mutants, could provide more space per bound DAXX molecule because these can spread out at the ends of the SPOP oligomers. The associated lower DAXX concentration in the resulting dense phase would be energetically favorable and could therefore explain why the saturation concentration of substrate with the interface mutants is reduced compared to SPOP^WT^.

Our proposed mechanism shows how linear SPOP self-association can have a non-monotonic effect on DAXX ubiquitination due to the emergent characteristics of SPOP-DAXX phase separation (43, 44). Briefly, some degree of self-association is necessary to amplify weak DAXX-DAXX interactions, but excessive self-association inhibits these same DAXX-DAXX interactions. Recent work on the mechanism of phase separation of Ubiquilin2 (UBQLN2) with poly-ubiquitin chains revealed the same physical principle (45). The soluble polyubiquitin “hub” binds and bundles UBQLN2, which has an intrinsic driving force for phase separation, thereby enhancing phase separation. However, concentrating UBQLN2 beyond its intrinsic dense phase concentration results in an energetic tradeoff. Poly-ubiquitin chains with different compactness therefore have different abilities to potentiate UBQLN2 phase separation, and an intermediate compactness seems to be ideal. Such tradeoffs between higher sticker density and optimal spacer length are expected to play roles in phase-separating systems in general.

The SPOP interface mutations characterized here were identified in patients with endometrial, skin, and other cancers, and their shared molecular mechanism was not previously understood. The majority of SPOP mutations identified in endometrial cancer are found at the periphery of the MATH domain (32), and the molecular mechanism underlying the resultant change in substrate specificity is not understood. Our conclusion that endometrial– and other cancer–related interface mutants alter SPOP phase separation behavior suggests that endometrial cancer–related MATH domain mutants also display altered phase behavior. Future research should assess this possibility.

Our work suggests that tuning of SPOP oligomerization could be used by the cell to modulate substrate levels. We tested this hypothesis by fusing GST dimerization domains to the SPOP N-terminus. While SPOP self-association was indeed enhanced and phase separation was reduced *in vitro*, the fusion proteins had very low expression levels in cells, and their activity could not be assessed. Oligomerization could be further tuned by post-translational modifications or by titration of the paralog SPOPL, a substrate recognition subunit that is highly homologous to SPOP, can dimerize with SPOP via its BTB domain, but has a dimerization-incompetent BACK domain due to an 18-residue insertion (Fig. 2C) and thus caps SPOP oligomers (30). The results we present here predict that varying SPOPL levels in cells may change the driving force for phase separation with certain substrates, and that this would alter their cellular levels. Given the tumor suppressor role of SPOP, future research into the regulation of its activity, including via modulation of its oligomer size distribution, will be valuable.

Our work also suggests a strategy for disentangling the contributions of condensates vs clusters in other multivalent systems. First, it is useful to recognize that condensates and clusters compete with each other; factors that stabilize clusters raise the saturation concentration. Second, while phase separation and cluster formation both use the same multivalent interactions (or stickers) for networking, phase separation requires an additional density transition, which is driven by the modest solubility of constituent monomers or complexes (27, 28). The solubility is determined by many properties of the protein including the surface charge distribution and conformation, meaning that both stickers and spacers contribute directly to protein solubility. Third, the network structures can differ between clusters and condensates (24). Hence, separation-of-function mutations to distinguish between the contribution from condensates and clusters should be accessible, either by modulating the solubility of molecules/complexes, or by differentially affecting the network structure of clusters vs condensates. A mutational strategy that capitalizes on these insights thus promises to reveal the extent to which function is mediated by phase separation in biology.

## Acknowledgments

T.M. acknowledges funding from NIH grant R01GM112846, the St. Jude Children’s Research Hospital Research Collaborative on The Biology and Biophysics of RNP Granules, and the American Lebanese Syrian Associated Charities. The content is solely the responsibility of the authors and does not necessarily represent the official views of the National Institutes of Health. The Ub-His and BRD3-Flag plasmids were kind gifts from Wenyi Wei (Harvard Medical School) and Joel Mackay (University of Sydney), respectively.

## Conflicts of Interest

T.M. was a consultant for Faze Medicines, Inc.

## SI Figures

**Figure S1.**
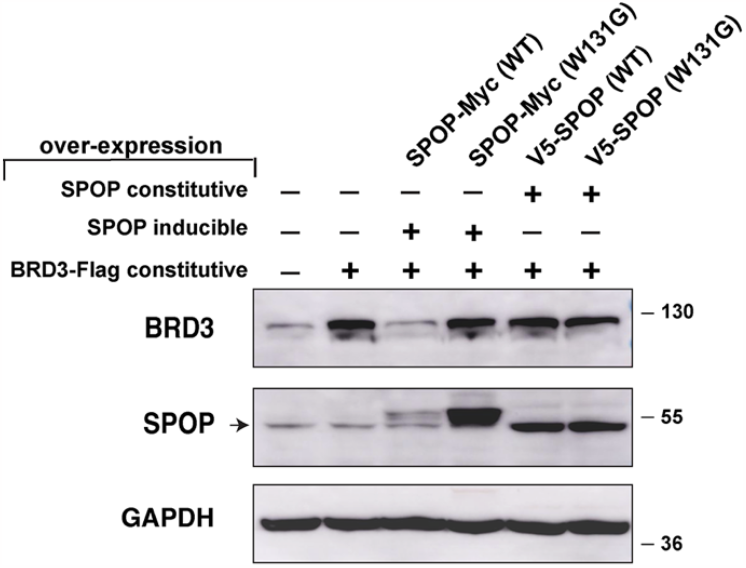
SPOP expression induced close to endogenous levels mediates BRD3 turnover. T-REx cells were transfected with plasmids for BRD3-Flag and SPOP-Myc (inducible expression) or V5-SPOP (constitutive expression). Twenty-four hours post-transfection, cells were lysed, and the resulting lysates were immunoblotted using antibodies for BRD3, SPOP, and GAPDH (loading control). The expected difference in BRD3 turnover for cells expressing SPOP^WT^ and SPOP^W131G^ is more pronounced with the induced-expression SPOP constructs.

**Figure S2.**
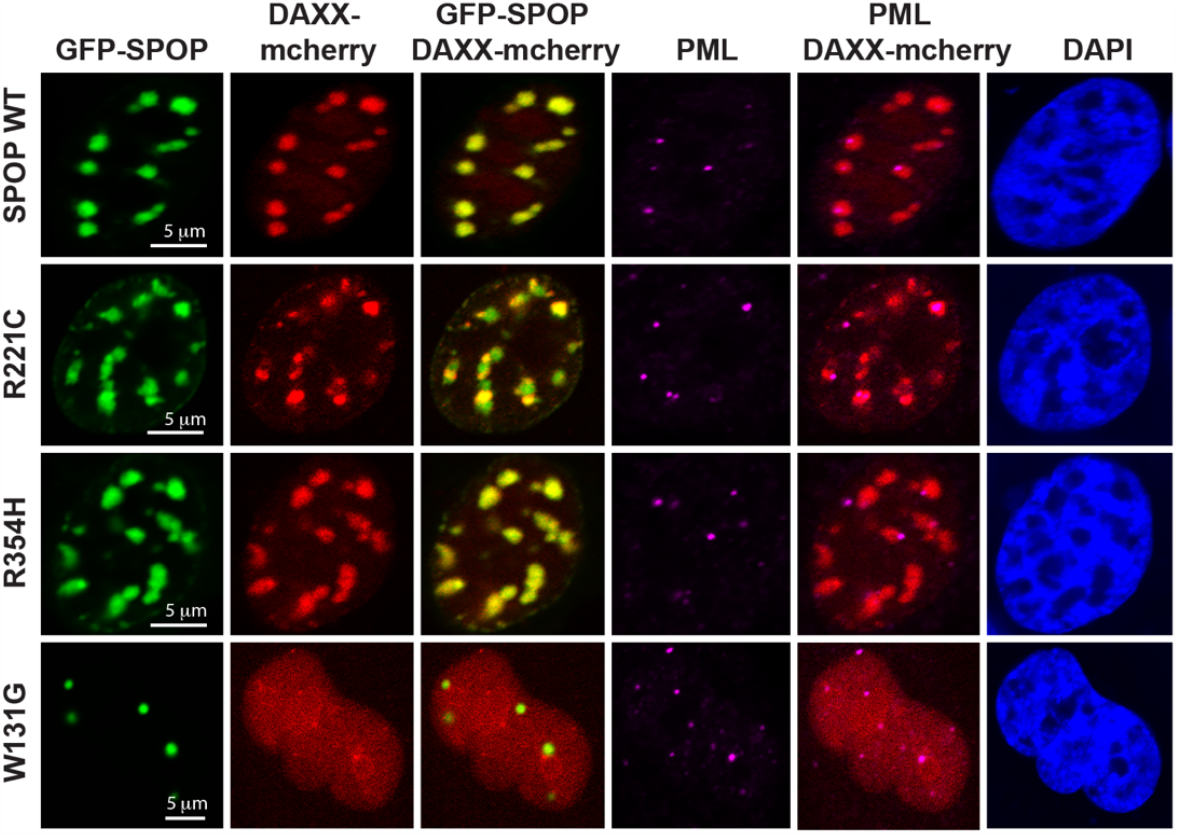
Representative fluorescence confocal images of HeLa cells expressing GFP-SPOP and DAXX-mCherry constructs. Cells were transfected with the indicated plasmids. Twenty-four hours post-transfection, cells were fixed and immuno-stained using an antibody against PML bodies (magenta). DAPI (blue) marks nuclear DNA.

## Notes

### Summary of Updates

A new citation to related work from another group has been added to the Discussion.

